# Using directed attenuation to enhance vaccine immunity

**DOI:** 10.1101/2020.03.22.002188

**Authors:** Rustom Antia, Hasan Ahmed, James J Bull

## Abstract

Many viral infections can be prevented by immunizing with live, attenuated vaccines. Early methods of attenuation were hit-and-miss, and these are now much improved by genetic engineering. But even current attenuation methods operate on the principle of genetic harm, reducing the virus’s ability to grow, which in turn limits the host immune response below that of infection by wild-type. We use mathematical models of the dynamics of virus and its control by innate and adaptive immunity to explore the tradeoff between attenuation of virus growth and the generation of immunity. Our analysis suggests that directed attenuation that disables key viral defenses against the host immune responses may attenuate viral growth without compromising, and potentially even enhancing the generation of immunity. We explore which immune evasion pathways should be attenuated and how attenuating multiple pathways could lead to robust attenuation of pathology with enhancement of immunity.

**Author summary:** Live attenuated virus vaccines are among the most effective interventions to combat viral infections. Historically, the mechanism of attenuation has been one of genetically reduced viral growth rate, often achieved by adapting the virus to grow in a novel cells. More recent attenuation methods use genetic engineering but also are thought to impair viral growth rate. These classical attenuations typically result in a tradeoff whereby attenuation depresses the within-host viral load and pathology (which is beneficial to vaccine design), but reduces immunity (which is not beneficial). We use models to explore ways of directing the attenuation of a virus to avoid this tradeoff. We show that *directed attenuation* by interfering with (some) viral immune-evasion pathways can yield a mild infection but elicit higher levels of immunity than of the wild-type virus.

## 1 Introduction

Many highly successful viral vaccines use live attenuated viruses, which are typically variants of the wild-type virus that have a genetically reduced capacity to grow [1, 2, 3]. Attenuated vaccines are considered to offer superior protection over other types of vaccines [4]. Early methods of attenuation were haphazard, relying on adaptation to viral growth in unnatural, artificial conditions to indirectly evolve a virus with reduced capacity to grow in the natural host. The fact that attenuation by this largely blind method was repeatable at all suggests that there is a broad window of viral growth reduction compatible with practical vaccine attenuation.

Viral attenuation entered a new era with genetic engineering, when it became possible to achieve quantitative reductions in viral growth rate and to block evolutionary reversion of attenuation [5, 6, 7, 8, 9, 10]. Much of the guesswork in reducing viral growth rate is eliminated with genetic engineering approaches, and the engineering can likewise slow or block the vaccine’s ability to reverse the changes and become virulent. Now, genetic engineering may be ushering in a third era, one in which attenuation is designed to enhance the immune response. Such ‘directed attenuation’ will require genetically altering (engineering) the virus in a manner that reduces pathology while enhancing the level of protective immunity. In this paper we use the term directed attenuation to refer to engineering viral genomes to influence the level of immunity.

The need for new methods of viral attenuation is not necessarily obvious. Many attenuated vaccines in use – created by old methods – have brought about dramatic reductions in the prevalence of infections with few side effects, so is there anything to be gained from changing their design? However, live attenuated vaccines typically do not elicit as much immunity as natural infection [11, 12, 13, 14, 15, 16]. This can potentially result in reinfection with circulating virus (e.g. for mumps where the and necessitates periodic boosting[12, 17]. Furthermore at a more fundamental level, vaccines remain elusive for many viruses [18, 19, 20, 21, 22] and require exploration of alternative approaches to vaccine design.

It is not a trivial matter to engineer a live vaccine and knowingly vary the immune response. It has become practical to engineer a virus to knowingly reduce the pathology (i.e., to attenuate, [4]). However current attenuation compromises the growth of the virus, leads to reduced viremia that typically elicits a smaller immune response [23]. The issue here is to attenuate the growth of the virus without compromising long term immunity, perhaps even elevating it. Doing so requires understanding the complex interactions and feedbacks between virus growth and generation of adaptive immunity.

The first step involves characterizing the different viral proteins and how they interact with the cells and molecules involved in the generation of immune responses – the subject of recent advances in molecular and cellular virology and immunology (reviewed in [24, 25]). Yet a qualitative understanding of these pathways is not enough: non-linear feedbacks between virus growth and immunity make it hard to comprehend the effect of modifying viral genes and proteins on the level of immunity generated. Can directed attenuation enhance immunity without increasing pathology? It is not clear if this is possible – targeting pathways that interfere with host immunity may simply result in more rapid virus clearance (less pathology) and consequently less clonal expansion and immunity at the end of the infection.

The approach to directed attenuation presented here is to develop a mathematical model that incorporates key elements of our current understanding the replicating virus and the immune response it elicits. We then use the model to explore whether directed attenuation might simultaneously reduce pathology and enhance immunity, and if so, to identify candidate pathways for virus engineering.

## 2 Model

We use a deliberately simple model for the dynamics of infection with virus or live-attenuated vaccine. Let *V* equal the virus density and *Z* and *X* equal the magnitude of the innate and adaptive responses respectively; all are functions of time.

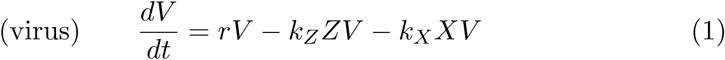

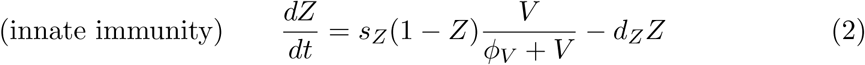

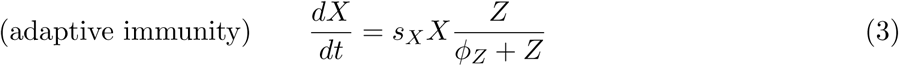

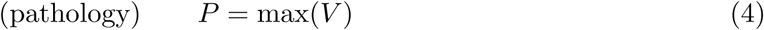

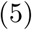

The virus grows exponentially at rate *r* in the absence of innate and adaptive immunity. Virus is killed by innate and adaptive immunity at rates proportional to current immunity levels and the parameters *k*_*Z*_ and *k*_*X*_ respectively. Innate immunity, *Z*, is activated by infection and saturates at 1, so *Z* equals the fraction of the the maximum value of innate immunity. Adaptive immunity, *X*, grows in proportion to innate immunity (*Z*). This representation is a modification of that usually used, where the stimulation of adaptive immunity depends only on the amount of virus *V*. It is justified on biological grounds that the generation of immune responses requires stimulation by innate immunity and that this can occur even after replicating antigen is cleared. We let the extent of pathology, *P*, equal the maximum level of virus. The basic form is similar to that used previously [26, 27, 28, 29, 30, 31].

The model is intended to capture basic properties of the immune response, helping discover unintuitive interactions and feedback effects that arise as a consequence of viral growth that interferes with the host’s immune response. This model does not pretend to capture all the known complexity of mammalian immune systems. For example, it does not include different populations of innate and immune cells or cytokines generated during immune responses. The inclusion of these details at this stage would lead to a complex model with no hope of parameterization. In these situations, simple models are often more robust and perform better than more complex ones [32, 33].

## 3 Results

### 3.1 Dynamics of acute infection

Dynamics of an acute infection are straightforward (Fig 1, showing changes in viral titer, innate and adaptive immunity over the course of infection). The virus initially grows exponentially. Soon thereafter the innate immune system is triggered and begins to limit viral growth. But innate immunity is temporary and decays to zero after the virus is cleared. The adaptive immune response is somewhat delayed relative to innate immunity, it is responsible for clearance of the infection and is subsequently maintained. As we are focused on the generation of adaptive immunity to the virus we do not consider subsequent waning of immunity at this stage.

**Figure 1:**
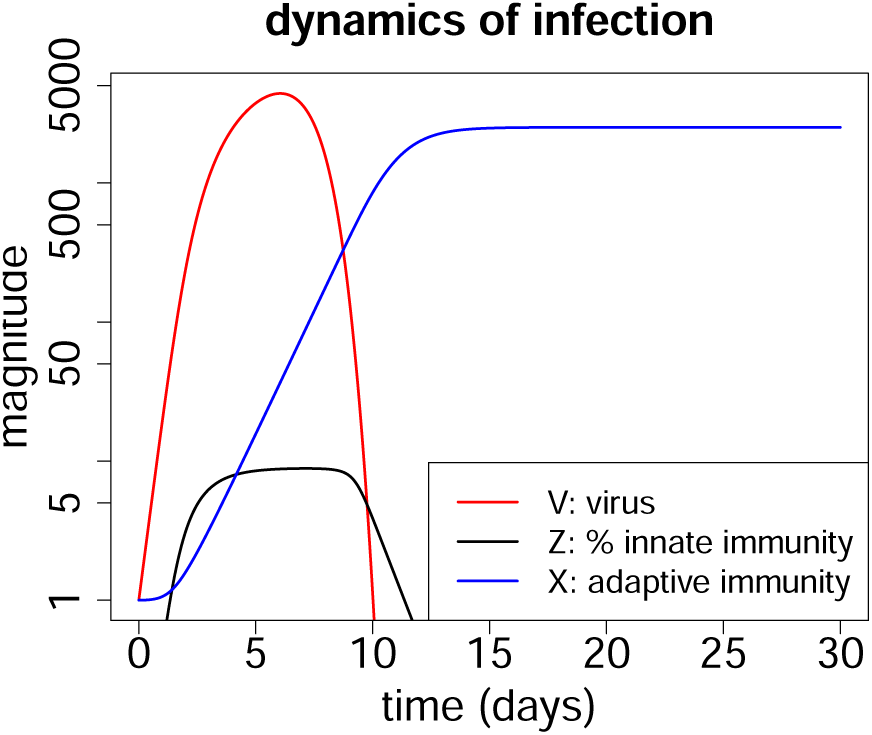
Typical dynamics of an acute infection. Virus (*V*) is shown in red, innate immunity (*Z*) in black and adaptive immunity (*X*) in blue. The scale for virus and immunity is fold change over the initial value (*V* (0), and *X*(0) are set to 1). The scale for innate immunity is percent of its maximum possible value, and in this simulation *Z* attains only about 20% of its maximum. Parameters are chosen for a biologically relevant regime as described earlier [34] and are shown in Table 1.

### 3.2 Classic attenuation by reducing virus growth

The original approach for attenuating a virus was to grow it in unnatural conditions (e.g., novel host or cell culture). Adaptation to the unnatural conditions often – but unpredictably – reduced growth rate in the primary host (humans), with consequent loss of pathogenesis. Newer methods of engineered attenuation also lower viral growth rate but do so far more predictably [4, 10]. With a viral growth rate less than wild-type, the attenuated virus attains in a smaller peak density before clearance and thus elicits less adaptive immunity. To wit, a single infection with wild-type measles, mumps or rubella viruses typically induces lifelong immunity [35, 36], while vaccine-induced immunity frequently requires boosting, as indicated by the CDC immunization schedule [17].

**Table 1:** Parameters of the model and the values used in simulations.

| Parameter | Description | Value and range |
| --- | --- | --- |
| $r$ | virus growth rate | $3\text{d}^{-1} (2 - 4)$ |
|  | <u>Innate immunity</u> |  |
| $s_Z$ | rate of activation of innate immunity | $.1\text{d}^{-1} (.06 - .14)$ |
| $\phi_V$ | sensitivity of innate immunity | $10^2\text{v} (10^1 - 10^3)$ |
| $d_Z$ | waning of innate immunity | $1\text{d}^{-1} (.4 - 1.6)$ |
| $k_Z$ | killing of virus by innate immunity | $30\text{vd}^{-1} (10 - 100)$ |
|  | <u>Adaptive immunity</u> |  |
| $s_X$ | rate of activation of adaptive immunity | $1\text{d}^{-1} (.5 - 1.5)$ |
| $\phi_Z$ | sensitivity of adaptive immunity | $.02\text{z} (.002 - .2)$ |
| $k_Z$ | killing of virus by adaptive immunity | $10^{-2}\text{xd}^{-1} (10^{-3} - 10^{-1})$ |

This basic trade-off provides a baseline that should be reproduced by any reasonable model of viral-immune dynamics: reduced viral growth rate should result in reduced viral density before clearance (reduced pathogenesis). There should be a consequent reduced stimulation of adaptive immunity and a lower final level of adaptive immunity. Our model indeed generates the expected patterns (Fig. 2). This pattern highlights the fundamental question of our study: is it possible to engineer an attenuation that is better than achieved by merely reducing viral growth rate – can we arrange pathogenesis to go down but immunity go up? Our approach to this question involves varying virus-effected parameters and observing changes in pathogenesis and in immunity, as done next.

**Figure 2:**
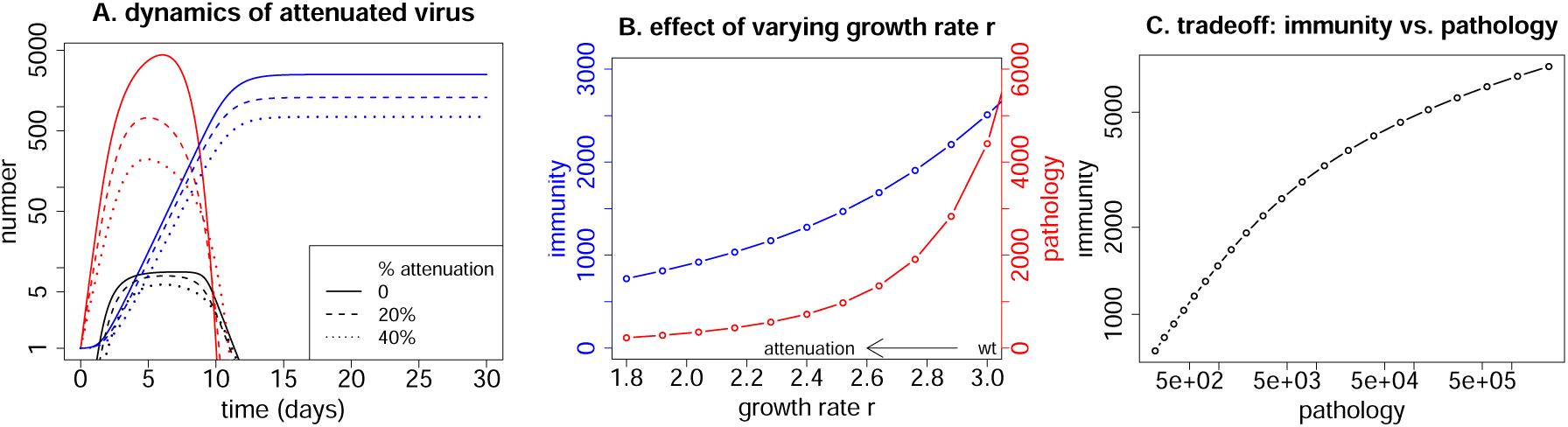
Attenuation by reducing the growth rate *r* of the virus. Solid lines indicate wild-type; dashed and dotted indicate attenuated. Reducing virus growth rate results in lower viremia as well as a reduction in the level of immunity. (A) Dynamics for the wild-type infection in solid lines (red for virus and blue for immunity) and for viruses with a 20% (dashed) and 40% reduced growth rate (dotted). (B) Impact of the degree of attenuation (reduction in *r*) on both the final level of immunity (blue) and the pathology (maximum virus density, red). (C) The tradeoff between pathology and immunity from changing growth rate *r*: reducing the growth rate results in lower pathology but also lower immunity. Parameters in Table 1.

### 3.3 Attenuation by suppressing virus immune evasion pathways

Viruses have pathways that interfere with the innate or adaptive immune responses [37, 38, 39, 40, 41]. As these reduce the magnitude or effect of immunity they are candidate pathways, that if targeted, could lead to an increase in immunity. We model the suppression of these viral pathways as changes in the parameters that correspond to attenuation strategies – reducing pathogenesis (Table 2). Note that suppression of a viral pathway may result in an increase of the parameter value – which can occur when the wild-type virus depresses a host anti-viral response.

**Table 2:** Candidates for directed attenuation, and summary of directed attenuation arising from changes in a single parameter. Desired outcomes that result in a decrease in pathology and an increase in immunity are shown in boldface.

| Evasion pathway targeted<br>(parameter & direction of change for vaccine) | Vaccine outcomes |  |
| --- | --- | --- |
|  | pathology | immunity |
| <u>Innate immunity</u> |  |  |
| max. activation rate for innate immunity ( $s_Z \uparrow$ ) | decrease | decrease |
| sensitivity of innate immunity to antigen ( $\phi_V \downarrow$ ) | decrease | no change |
| **decay of innate immunity** ( $d_Z \downarrow$ ) | **decrease** | **increase** |
| killing by innate immunity ( $k_Z \uparrow$ ) | decrease | decrease |
| <u>Adaptive immunity</u> |  |  |
| **max. rate of clonal expansion** ( $s_X \uparrow$ ) | **decrease** | **increase** |
| **sensitivity of immunity to antigen** ( $\phi_Z \downarrow$ ) | **decrease** | **increase** |
| killing by adaptive immunity ( $k_X \uparrow$ ) | decrease | decrease |

Virus strategies to evade innate immunity are modeled as low values of *s*_*Z*_ (the maximum rate of stimulation), high values of *ϕ*_*V*_ (the level of virus needed for stimulation) and high values of *d*_*Z*_ (the decay rate of innate immunity). Similarly viruses avoid stimulation of adaptive immunity by low values of *s*_*X*_ (the maximum rate of proliferation of immune cells) and high values of *ϕ*_*Z*_ (the level of stimulation required to induce proliferation of adaptive immune cells). Viruses can also limit killing by the innate and adaptive immunity with low values of the rate constants *k*_*Z*_ and *k*_*X*_ respectively. Attenuation thus involves changing those parameters in the opposite directions, reducing pathology. But our interest also lies in which of these changes will have the additional effect of increasing the final level of adaptive immunity, or at least not lowering it.

#### Blocking immune-evasion genes attenuates the infection; some changes can also enhance immunity

Blocking (reducing) any viral immune-evasion pathway leads to attenuation of the virus (Fig 3). All are thus potential routes for generation of a live attenuated virus vaccine. But how might those changes affect the level of immunity? Blocking pathways that make the virus more sensitive to clearance by either innate (*k*_*Z*_) or adaptive (*k*_*X*_) immunity results in more rapid control of the virus but a lower final level of immunity. In contrast, blocking pathways that slow the generation of adaptive immunity (either by changing the immune growth rate, *s*_*X*_, or sensitivity of adaptive immunity, *ϕ*_*Z*_) results in greater stimulation and a *higher* final level of adaptive immunity. Finally, suppressing the viral pathway accelerating innate immune decay (*d*_*Z*_) prolongs the stimulation of adaptive immunity and increases its final level.

**Figure 3:**
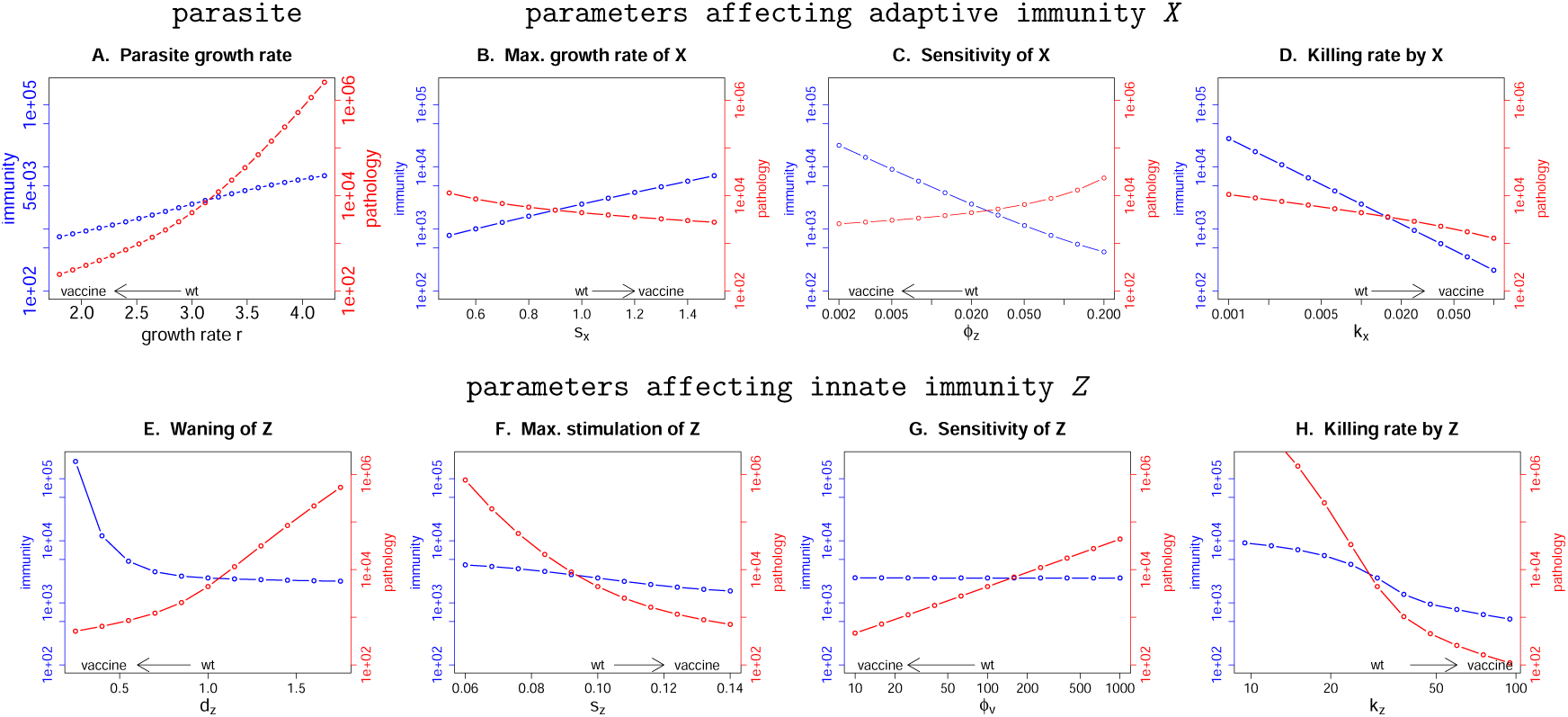
Effect of changing single parameters on the levels of pathology, immunity. Red curves give the pathology, with scale on the right vertical axis; blue lines give the final level of immunity, with scale on the left vertical axis. The baseline parameters chosen for the wild-type virus are indicated by ‘wt’ on the horizontal axis. A vaccine strain would be designed to lower pathology, and the arrow immediately above the horizontal axis gives the direction of change in the parameter value that would reduce pathology. Baseline parameters are as in Fig 1 except for the parameter whose value is changed in the panel.

Thus attenuation by modifying immune-evasion pathways described by parameters *s*_*X*_, *ϕ*_*Z*_ and *d*_*Z*_ generates the hoped-for outcome of reduced pathology with increased immunity. Attenuation of other immune-evasion pathways, such as those associated with changes in the stimulation of innate immunity (*s*_*Z*_ or *ϕ*_*V*_) or susceptibility to killing by innate or adaptive immunity (*k*_*Z*_ and *k*_*X*_), does not result in higher immunity. A more detailed consideration of the effect of changing parameters on the dynamics of infection and immunity is considered in supplemental file S1.

#### A pathology-immunity plot enables easy comparison of attenuation strategies

The effects of different attenuation pathways may be directly compared by plotting them together in a grid of pathology and immunity (Fig 4). The ideal live attenuated virus vaccine would generate lower pathology but higher immunity than infection with the wild-type virus, corresponding to the upper left quadrant (where wild-type is taken as the center). It is straightforward to see that directed attenuation works for the same three parameters identified above, *d*_*Z*_, *s*_*X*_ and *ϕ*_*Z*_. It is also seen that increasing sensitivity of innate immunity (decreasing *ϕ*_*V*_) leads to attenuation of pathology without compromising immunity. A summary of successful directed attenuation strategies by single parameter changes is given in Table 2.

**Figure 4:**
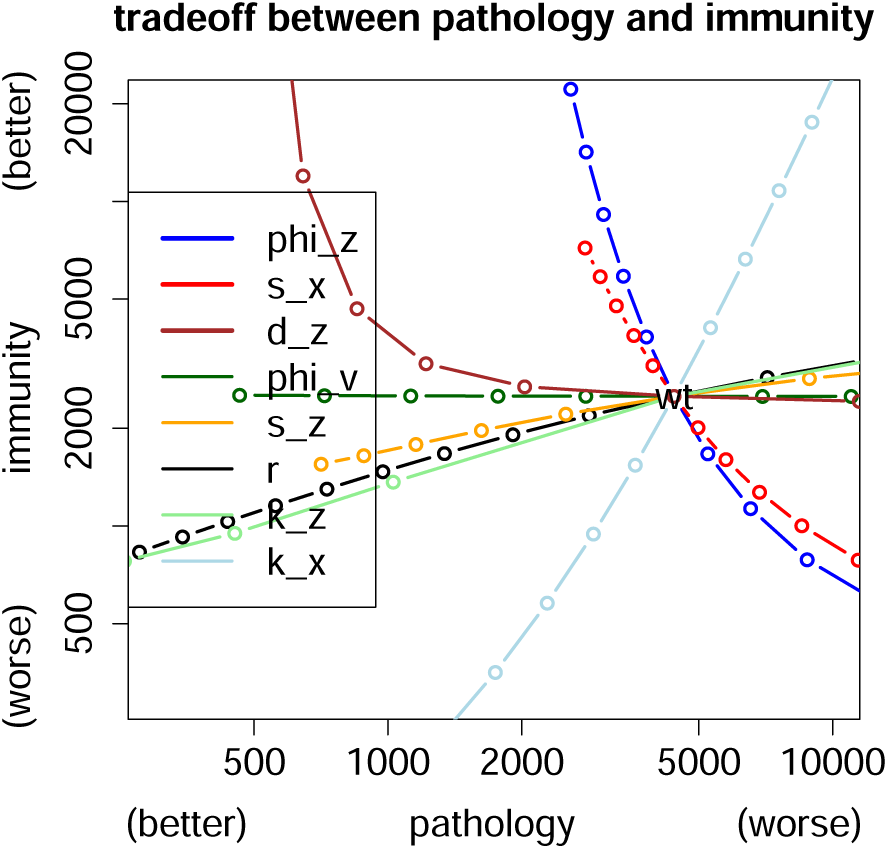
Comparison of immunity-pathology tradeoffs generated by changing single parameters. Wild-type values are given at the intersection of the curves, so viable attenuation strategies would lie to the left. As the goal is to attenuate and to increase the immune response, the desirable attenuation strategies lie in the upper left quadrant. The tradeoff for the classic mode of attenuation – lowering growth rate *r* (black line) – has the undesirable effect of lowering immunity. In contrast, decreasing the rate of loss of innate immunity (*d*_*Z*_ ↓), or increasing the rate of proliferation or sensitivity of adaptive immunity (*s*_*X*_ ↑ (red), *ϕ*_*Z*_ ↓ (blue)) leads to lowering pathology and increasing immunity.

#### Combining attenuation strategies

Two potential problems with directed attenuation of a single viral anti-immune pathway are reversion and the consequences of vaccination of immunocompromised individuals. As we discuss later reversion may not be a significant problem, and we focus on inadvertent vaccination of immunocompromised individuals. The problem is especially apparent if a virus is attenuated by deletion of an antiviral strategy that targets a defense pathway lacking in the patient. The classic method of attenuation which compromises the growth of the virus (reduces the growth rate *r*) avoids this problem, or at least makes the infection less severe than that with wild-type virus. However if an individual was To overcome this problem, directed attenuation could use a combination strategy. One type of combination is to block immune evasion while separately reducing growth rate. Another type of combination is to block both evasion of adaptive immunity and evasion of innate immunity.

A worry with combination strategies is that the reduced immunity might be too severe: can wild-type immunity be attained in a virus with classic growth rate reduction when an anti-immunity defense is also disabled? The answer appears to be affirmative for some combinations (Fig 5). The top four panels show the effect of reducing viral growth rate (going from right to left on the horizontal axis) with the effect of changing one immune parameter. Suppressing an immune-evasion pathway that changes the sensitivity of the adaptive immunity (reduces *ϕ*_*Z*_) or the decay rate innate immunity (i.e. increases *d*_*Z*_) restores vaccine immunity to the level elicited by the wild-type virus (right column), and this is done without compromising the level of attenuation of pathogenesis that was obtained by the reduction in *r* (left column).

**Figure 5:**
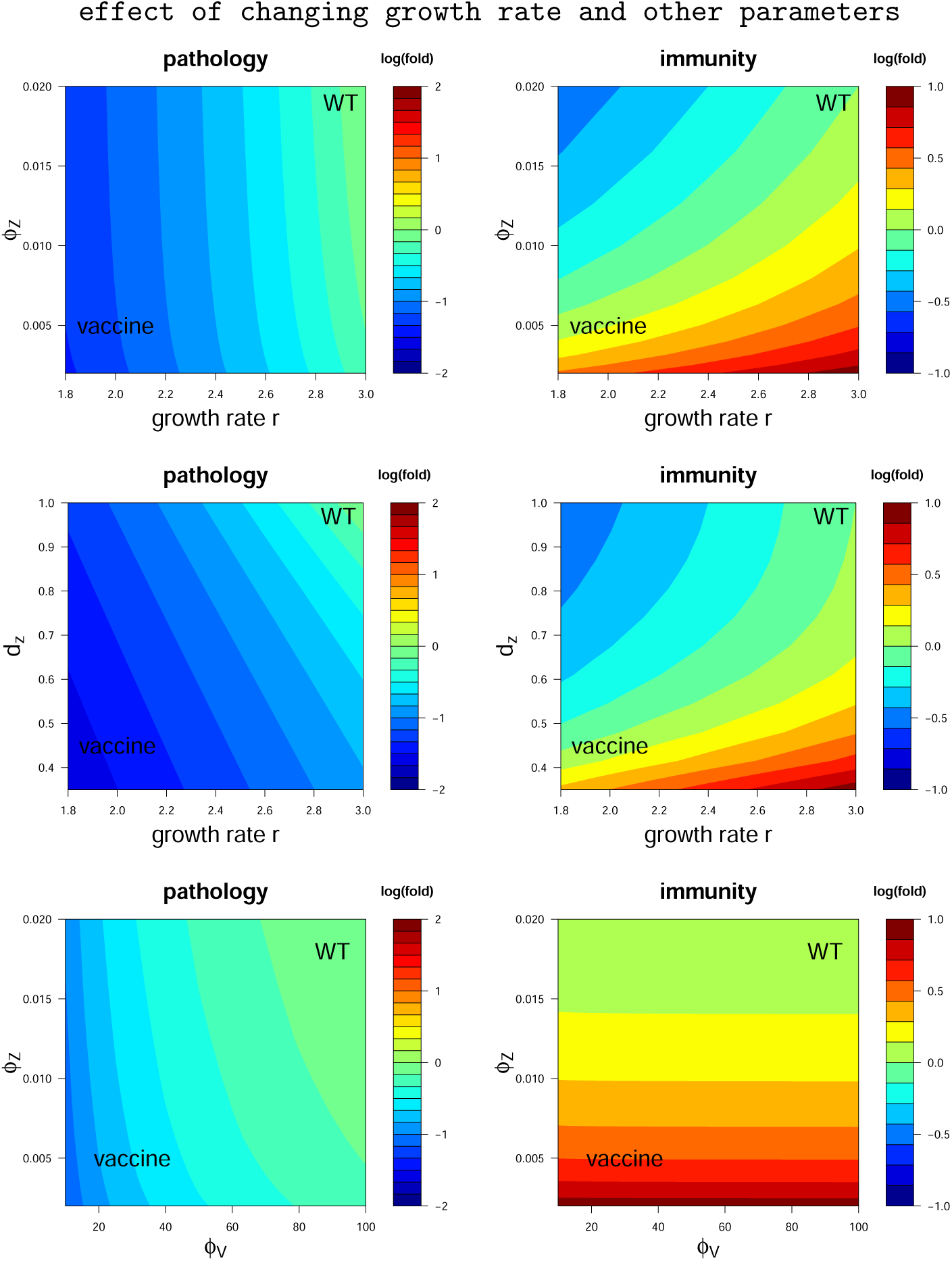
Heat maps for the log_10_ fold changes in pathology (left column) and immunity (right column) when modifying two parameters. The top two rows include growth rate as one of the parameters changed (horizontal axis) – the classical attenuation method. The effect of changing a single parameter alone is seen by moving parallel to the respective axis. The goal of attenuation is to reduce pathology from wild-type values (increase the level of blue, left column) and to increase immunity (increase redness, right column). Values of wild-type virus are given in upper right of each panel, values of the prospective vaccine in lower left. The goal is to have pathology become increasingly blue and immunity become increasingly red in traversing from wild-type to vaccine. The classical attenuation strategy arising from reductions in the single parameter *r* is seen to reduce both pathology and immunity (moving left along the horizontal axis in any of the top four panels). However, combining reductions in *r* with increasing the sensitivity of the adaptive immune response (i.e. decreases *ϕ*_*Z*_) or increasing the duration of innate immunity (i.e. increases *d*_*Z*_) restores the level of immunity generated by the vaccine to that induced by the wild-type virus (top two rows). The bottom row shows that ideal attenuation can be achieved by changes in a pair of parameters that does not include *r*.

Combinations are possible without altering growth rate (*r*), and some combinations may likewise result in reduced pathology with enhanced immunity (Fig. 5). The combination shown that does not change growth rate (bottom row) suppresses viral evasion of innate immunity (increasing *ϕ*_*V*_) while sensitizing adaptive immunity to stimulation by innate immunity (increasing *ϕ*_*Z*_).

One of the striking observations from Fig. (5) is that the effect of varying parameters is substantially different for the 3 cases. In the top row, immunity changes in response to both parameters, but pathology changes largely in response to the growth rate *r*. In the bottom row, immunity is strongly affected by the sensitivity of the adaptive response *ϕ*_*Z*_ but largely unaffected by the sensitivity of the innate immune response *ϕ*_*V*_, whereas pathology shows much the opposite pattern. The middle row shows both pathology and immunity being affected by the combination of parameter values. These results reinforce the unintuitive nature of directed attenuation, and illustrate how models can be useful tools to understand the consequences of the non-linearities and feedbacks for different vaccination strategies.

## 4 Discussion

Attenuated vaccines have been the mainstay of viral vaccines for close to a century [2, 3, 4]. Attenuation is typically interpreted as a genetic reduction in viral growth rate (reduced ability to grow in the host), an outcome that should be easily achieved by almost any disruptive modification to the viral genome. Advances in genetic engineering now enable attenuated designs that were previously unattainable, but even these have tended to be of a mode that disrupts viral growth [5, 6, 7, 8, 9, 10].

Here we suggest an extended approach to attenuation, one that directs the virus toward reduced defenses against host immunity. This approach to attenuation is suggested by the fact that many viruses directly interfere with the immune response [37, 38, 42, 43, 39, 40, 41], and those interference genes are obvious targets for genetic engineering. Targeting viral immune-evasion pathways compromises the viral load and in general results in a reduction of pathology. However, unlike reducing the virus growth rate which not only reduces pathology but also reduces the immune response, ‘directed’ attenuation might instead increase the immune response. Using a simple computational model of the immune response to viral infection, we indeed find that disruptions of some, but not all, viral anti-immune responses can attain the desirable goal of reduced pathology without compromising the generation of immunity. Our model allows us to identify characteristics of the immune-evasion pathways that should be targeted to achieve the desired outcome. Changing more than one parameter in the model allows us to explore the consequences of targeting multiple virus pathways. We suggest that an attenuated design that includes reduced viral growth rate together with disabling carefully chosen immune-evasion pathways would be ideal, in that it could provide enhanced immunity compared with natural infection along with reduced pathology, even in immune-compromised individuals.

While we have focused on virus pathways that compromise the magnitude of the immune response, similar principles could be applied to virus pathways that compromise the maintenance of memory. It could also be extended to inactivated virus vaccines where genetic engineering to remove genes for virus proteins that inhibit the generation of immune responses may (if the proteins function in the absence of virus replication) enhance the immune response elicited by vaccination with the killed virus.

### Examples of immune evasion strategies

Recent studies have discovered many pathways used by viruses to evade the innate and adaptive immune responses [37, 38, 39, 40, 43, 41]. Poxviruses are large DNA viruses with much of their genome encoding immune-evasion pathways [39, 40, 44]. These pathways target host type 1 interferon, tumor necrosis factors and the complement pathway of innate immunity. They also evade adaptive immunity by downregulating antigen presentation and by blocking costimulatory pathways and the apoptotic response. Adenoviruses are medium-sized, non-enveloped viruses containing a double stranded DNA genome. They encode immune-evasion pathways that inhibit tumor necrosis factor activity, and also evade adaptive immunity by downregulating antigen presentation [45, 46, 47].

One of the best studied immune-evasion genes is the NS1 protein of the influenza virus. The major function of NS1 protein is to interfere with multiple stages of the type 1 interferon signaling cascade [48, 49, 50, 51, 52]. Mutant influenza virus lacking NS1 protein are highly attenuated in wild-type (interferon-competent) mice, but not in IFN-incompetant systems such as STAT 1 knockout mice [53]. Viruses with a truncation or deletion of their NS1 gene have been shown to be promising candidates for a live attenuated vaccine in chicken [54, 55]. Type 1 interferon plays a role in reducing virus replication, which corresponds to a decrease in the growth rate of the virus (*r*) in our models without changing parameters that govern the generation of immune responses. Changes in only the parameter *r* in our model results in less pathology and a smaller immune response. This is consistent with the outcome of experimental infections of chicken with influenza virus vaccines that have deletions in the NS1 protein [55]. We suggest that it might be worth exploring the possibility of targeting other virus immune-evasion pathways so that the immune response is increased.

### How to direct attenuation

Our results are based on a simple model of the within-host dynamics of virus and immunity during acute infections. Applying these models to specific viruses will require incorporating details of how that particular virus evades the immune response, as well as the relevant components of the immune response elicited by the virus.

Selecting appropriate immune-evasion pathways will certainly require experimental studies. These experiments would be along the lines of the studies of NS1 deletion of the influenza virus discussed earlier. The first step would be the generation of potential vaccine viruses by attenuating different immune-evasion genes, ideally in a graded manner. The vaccine candidates would be used to infect animals, allowing quantification of how different attenuation strategies affect both the dynamics of virus (and pathology) and the generation of the immune response. Intuiting the consequences of different mutations is challenging because of the non-linear feedbacks between control of virus and generation of host immunity. Consequently, models that incorporate the relevant details of specific virus immune-evasion pathways may help suggest combinations of pathways to be targeted, and analysis of the results will in turn allow refinement of the models.

At a more basic engineering level, we suggest that in addition to compromising virus immune-evasion genes (e.g., knockouts or codon deoptimizations), directed attenuation can involve up-regulation or addition of genes that enhance the generation of immunity. Candidates for up-regulation or addition include pathogen associated molecular patterns (PAMPS) that induce the appropriate form of immunity. Studies on surrogates of protection [56] could help identify appropriate PAMPS for inclusion in the virus.

One consideration is the possibility of virus reversion. Gene additions are more likely to be prone to reversion through deletions and point mutations than gene deletions [57]. However they (i.e. [57]) suggested that within-host vaccine evolution can compromise vaccine immunity only when the extent of evolution during vaccine manufacture is severe, and that this evolution may be avoided or mitigated by suitable choice of the vaccine inoculum.

### Caveats, limitations and future directions

We have intentionally used simple models to illustrate the potential for directed attenuation strategies for the development of vaccines. This is because the particularities of each virus and its evasion mechanisms would require a model tailored to the relevant details, and generating one complex and general model is both unrealistic and intractable at the current time. There are some empirical complexities to include when considering the relevant details for a particular virus. At least for viruses with small genomes, genes with anti-immune functions may have pleiotropic effects, and it may be challenging to engineer them to retain other potentially essential functions while disrupting their anti-immune functions. Pleiotropic effects resulting from disruption of an immune-evasion pathway could be incorporated into the models by allowing changes in more than one term and parameter. The current models would need to be extended to include scenarios where the extent of attenuation (which we describe by a change in the relevant parameters) depends on virus density, as has been suggested for some non-selfish virus evasion strategies [58]. Our current models of immunity do not include different branches (e.g., humoral and cellular) or the complex differentiation pathways for the generation of immunological memory. Elaborations of the model are warranted, but any elaborations should be based on detailed consideration of a vaccination scenario for a specific virus. Simple models omit many details but have the advantage that they are often more robust than highly complex ones [32, 33]; the current model incorporates the feedbacks between viral defenses and immune responses, which may be the key properties relevant to directed attenuation.

### Summary

This study uses models to explore the consequences of attenuating a virus by targeting the interaction of the virus with the host’s immune response, rather than by the more traditional approach of targeting viral growth rate. This approach allows us to explore how new vaccines may be designed with the goal of maximizing the level of protective immunity generated. Contrary to the conventional view, it should be possible to engineer vaccines that provide stronger protection than that provided by natural infection.

## 5 Supplementary Information

In the SI we use simulations of the dynamics of infection to give us an insight into the interactions that give rise to the tradeoffs observed when we attenuate various immune-evasion pathways of the virus.

### Dynamics of attenuation by modulation of innate immunity

Viruses can attenuate or evade innate immunity in a number of ways. In terms of the parameters of the model attenuation of innate immunity can occur by changes in 4 parameters, *s*_*Z*_ the rate at which innate immunity can be stimulated, *ϕ*_*V*_ the sensitivity of innate immunity to recognizing virus, *d*_*Z*_ the rate of waning of innate immunity, and *k*_*Z*_ the rate constant for killing of the virus by innate immunity.

The sensitivity of innate immunity can be increased by reducing the virus density at which innate immunity is triggered, *ϕ*_*V*_ or the rate at which innate immunity is stimulated *s*_*Z*_. As seen in Fig 3 In both these scenario’s we find that there is a significant reduction in the pathology but only a small reduction in the level of adaptive immunity generated. The reason for this can be seen in the representative simulations shown in Fig 6A&B. We see that the more rapid generation of an innate response results in more rapid control (but not clearance) of the virus by innate immunity. As the adaptive immune response is needed for virus clearance we find that the duration of infection is not significantly affected and consequently the amount of adaptive immunity generated does not substantially. Thus changing these two parameters can lead to a reduction in the level of pathology and smaller reduction in the generation of immunity.

**Figure 6:**
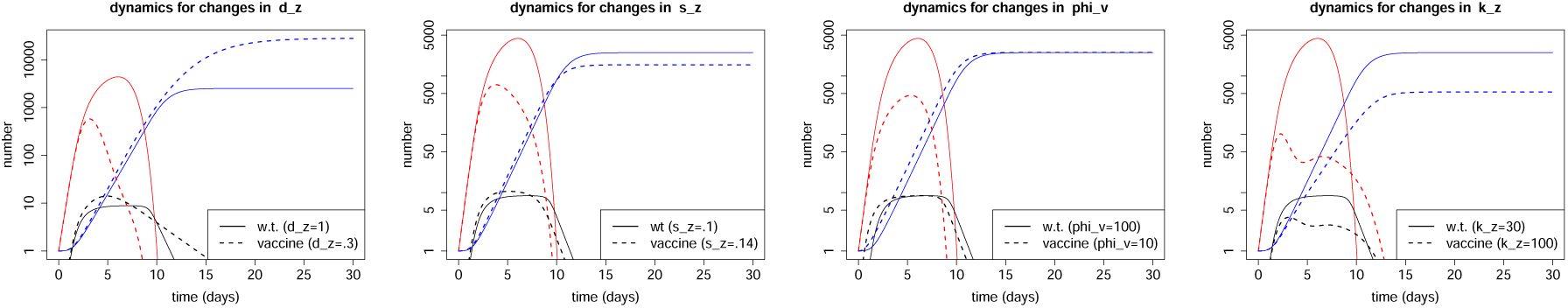
Dynamics of responses of vaccines that attenuate evasion of innate immunity. The plots show the dynamics of virus in red, innate and adaptive immunity in black and blue. The dynamics of the response to wild type virus is shown by the solid lines and the vaccine strain by the dashed lines. Parameters as in Fig 1 except as indicated.

Another way a virus can also evade innate immunity is by increasing the rate of decay of innate immunity, *d*_*Z*_. As seen in Fig 3, a vaccine strain of the virus that counters this by reducing the rate of decay of innate immunity will not only reduce the extent of pathology but interestingly can generate more immunity than is elicited by the wild-type vaccine strain. The cause of this can be seen in Fig 6C where we find that the reduction in *d*_*Z*_ results in a larger and longer innate response. This response controls the virus at a lower density and the longer duration of the response increases the stimulation of the adaptive immune response.

The final way which consider for evasion of innate immunity is by decreasing the rate at which innate immunity kills the virus. A vaccine strain of virus that has an increased susceptibility to innate immunity will have lower pathology and lower immunity (as seen in Fig 3). Fig 6D shows the cause – the virus is more rapidly controlled and this results in a lower level of innate immunity and less stimulation of adaptive immunity.

### Dynamics of attenuation by modulation of adaptive immunity

Viruses can attenuate or evade adaptive immunity in a number of ways. In terms of the parameters of the model attenuation of adaptive immunity can occur by changes in 3 parameters, *s*_*X*_ the maximum rate at growth of adaptive immunity, *ϕ*_*Z*_ the sensitivity of stimulation of adaptive immunity to recognizing virus and *d*_*Z*_ and *k*_*X*_ the rate constant for killing of the virus by adaptive immunity.

As seen in Fig 3 changes to the virus that increases in the rate of proliferation of adaptive immunity, either by increasing the rate of proliferation of immune cells *s*_*X*_ or increasing the sensitivity of immune cells to stimulation *ϕ*_*V*_ results in attenuation (less pathology) and higher levels of immunity. The reason for this can be seen in Fig 7 where we see that increases in *s*_*X*_ or decreases in *ϕ*_*V*_ result in faster / earlier growth of adaptive immunity. In both cases the more rapid generation of adaptive immunity results in earlier clearance of the pathogen and curtailing the both innate immunity and the stimulation of adaptive immunity – though the decrease in *ϕ*_*V*_ can to some extent compensate for this.

**Figure 7:**
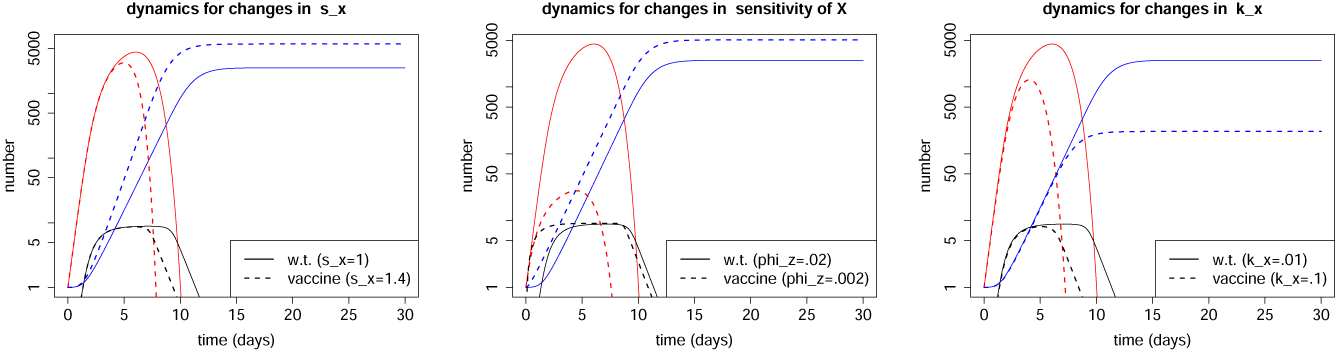
Dynamics of responses of viruses with enhanced levels of inducing and susceptibility to adaptive immunity. The plots show the dynamics of virus in red, innate and adaptive immunity in black and blue. The dynamics of the response to wild type virus is shown by the solid lines and the vaccine strain by the dashed lines. Parameters as in Fig 1 except as indicated.

As is the case for innate immunity, an increase in the rate of clearance of the virus by adaptive immunity leads to more rapid clearance of the virus which causes a decrease in pathology but also a decrease in the extent of stimulation and final level of adaptive immunity (see Fig 7).

